# TGF-β2, catalase activity, H_2_O_2_ output and metastatic potential of diverse types of tumour

**DOI:** 10.1101/467191

**Authors:** Malak Haidar, Mehdi Metheni, Frederic Batteux, Gordon Langsley

**Author notes:** co-first authors, equally contributed (Malak Haidar, Mehdi Metheni). Corresponding author: Gordon Langsley.

## Abstract

*Theileria annulata* is a protozoan parasite that infects and transforms bovine macrophages causing a myeloid-leukaemia-like disease called tropical theileriosis. TGF-β2 is highly expressed in many cancer cells and is significantly increased in *Theileria*-transformed macrophages, as are levels of Reactive Oxygen Species (ROS), notably H_2_O_2_. Here, we describe the interplay between TGF-β2 and ROS in cellular transformation. We show that TGF-β2 drives expression of *catalase* to reduce the amount of H_2_O_2_ produced by *T. annulata*-transformed bovine macrophages, as well as by human lung (A549) and colon cancer (HT-29) cell lines. *Theileria*-transformed macrophages attenuated for dissemination express less catalase and produce more H_2_O_2_, but regain both virulent migratory and matrigel traversal phenotypes when stimulated with TGF-β2, or catalase that reduce H_2_O_2_ output. Increased H_2_O_2_ output therefore, underpins the aggressive dissemination phenotype of diverse tumour cell types, but in contrast, too much H_2_O_2_ can dampen dissemination.

## Introduction

TGF-β (Transforming Growth Factor beta) is a pleiotropic cytokine that is involved in diverse cellular processes such as proliferation, apoptosis and motility (1, 2). In advanced cancer, TGF-β acts as an oncogenic factor that promotes tumour progression. Three isoforms have been defined of which TGF-β2 is highly expressed in many cancer cell lines, especially those showing a high dissemination potential. *Theileria annulata* parasitizes bovine macrophages and transforms them into disseminating tumours that cause a myeloid-leukemia-like disease called tropical theileriosis. However, *T. annulata*-transformed macrophage dissemination can be attenuated by multiple *in vitro* passages and attenuated macrophages are used as live vaccines in countries endemic for tropical theileriosis (3).

TGF-β2 levels are high in *Theileria*-transformed macrophages and this correlates with susceptibility to disease (4). Upon attenuation *in vitro*, the Ode vaccine line displays both reduced TGF-β2 expression and dissemination, which are re-established by addition of exogenous TGF-β (4); observations consistent with a pro-metastatic role for TGF-β2 in the virulence of *Theileria*-transformed macrophages. Furthermore, excessive cellular oxidative stress diminishes the virulence of *Theileria*-transformed macrophages, as attenuated macrophages produce more H_2_O_2_ (5). However, heightened reactive oxygen species (ROS) have been reported to increase TGF-β expression and stimulate release of TGF-β from latent complexes (6, 7). Although this might contribute to TGF-β2 production by virulent macrophages it does not explain why attenuated macrophages that produce more ROS express less TGF-β2 (4, 5). TGF-β2-signalling induces the transcription factor CREB in *Theileria*-transformed macrophages and CREB activity diminishes upon attenuation of macrophage virulence (8). The cyclic AMP response element-binding protein (CREB) is a transcription factor of general importance in diverse cell types (9). CREB signalling is associated with cancer development and poor clinical outcome in leukemogenesis (10), but it is not known if CREB is a player in ROS regulation.

Elevated rates of ROS production have been described for human cancer cells, where excessive ROS underpins many aspects of tumour development and progression (11). However, tumours also express increased levels of antioxidant proteins to detoxify ROS such as superoxide dismutases (SODs), catalase, peroxiredoxins, the glutathione system that includes glutathione (GSH), glutathione reductase and glutathione peroxidases (GPx) (11). Here, we focus on catalase that detoxifies hydrogen peroxide (H_2_O_2_) by turning it into water and oxygen. We report that TGF-β2 induces CREB transactivation to promote catalase transcription that leads to increased catalase activity and reduction in H_2_O_2_ levels. We provide evidence that TGF-β2-driven catalase activity regulates the H_2_O_2_ redox balance that impacts directly not only on the hyper-dissemination phenotype of *Theileria*-transformed macrophages, but also on the metastatic potential of human lung and colon cancer cell lines.

## Results

### *T. annulata*-transformed macrophages attenuated for dissemination display significantly less catalase activity compared to virulent hyper-invasive macrophages

We previously showed that attenuation of *Theileria*-transformed macrophage virulence correlates with an increase of H_2_O_2_ output (5). This appeared counter intuitive, as one imagined that a decrease in infected macrophage virulence would be accompanied by a reduction in their oxidative stress status. However, accumulation of H_2_O_2_ can occur either due to an increase in superoxide dismutase (SOD) activity that produces H_2_O_2_, or reduced detoxification of H_2_O_2_ leading to its accumulation. In order to discriminate between these two possibilities SOD and catalase activities were measured in virulent (V) and attenuated (A) *Theileria*-transformed macrophages (Figure 1). No significant change in SOD activity was detected between virulent and attenuated macrophages (Fig. 1A), whereas catalase activity was significant diminished in attenuated macrophages (Fig. 1B). Moreover, decreased catalase protein levels underpinned the reduced catalase activity of attenuated macrophages (Fig. 1C). Catalase activity was also measured in *Theileria*-transformed macrophages isolated from disease-resistant Sahiwal cattle and found to be lower (Fig. S1A). Thus, attenuated macrophages isolated from disease-susceptible animals resemble transformed macrophages isolated from disease-resistant animals with respect to catalase activity.

**Figure 1:**
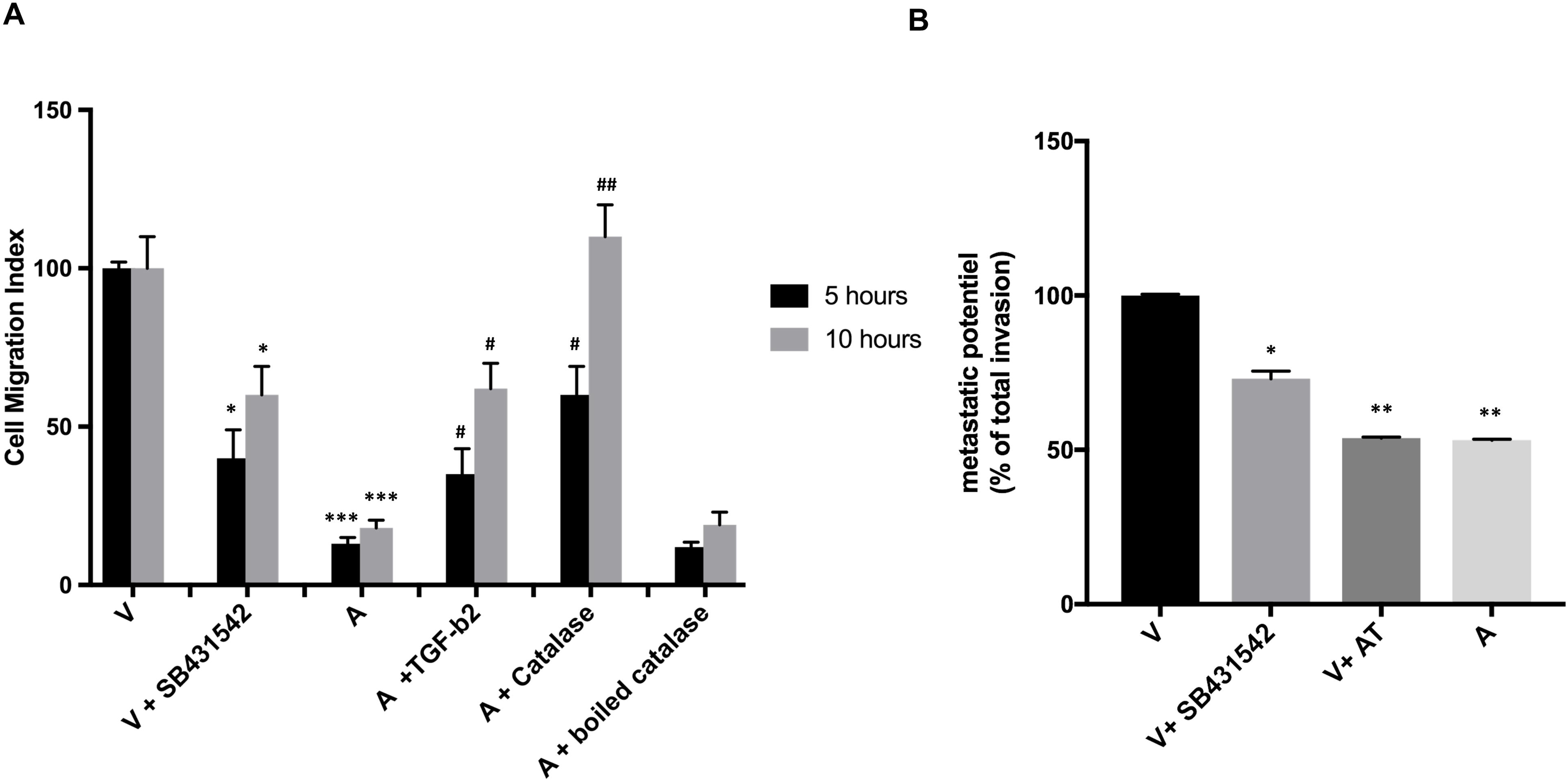
Catalase activity is decreased in attenuated *Theileria*-transformed macrophages. **A.** SOD activity doesn’t change between virulent and attenuated macrophages. **B.** Catalase activity is diminished in attenuated compared to virulent macrophages. **C top**, Catalase expression is high in virulent macrophages compared to attenuated macrophages. **C bottom**, Actin expression is unchanged between virulent and attenuated macrophages. ROS measurements in A and B were done independently (n = 3) and in triplicate. **p<0.005 compared to attenuated macrophages.

### TGFβ-2 stimulates *catalase* transcription leading to an increase in protein levels and catalase activity

We examined if the decrease in TGF-β2 levels underpinned loss of catalase transcription and activity. When attenuated macrophages were stimulated with recombinant TGF-β2 *catalase* transcription is increased, but not that of *glutathione peroxidase*, coding for another antioxidant enzyme (Figure 2A). Stimulation of attenuated macrophages with recombinant TGF-β2 increased catalase activity (Fig. 2B) and decreased H_2_O_2_ output (Fig. 2C). Taken together, TGF-β2-signalling clearly activates *catalase* transcription leading to increased amounts of catalase and greater catabolic activity towards H_2_O_2_.

**Figure 2:**
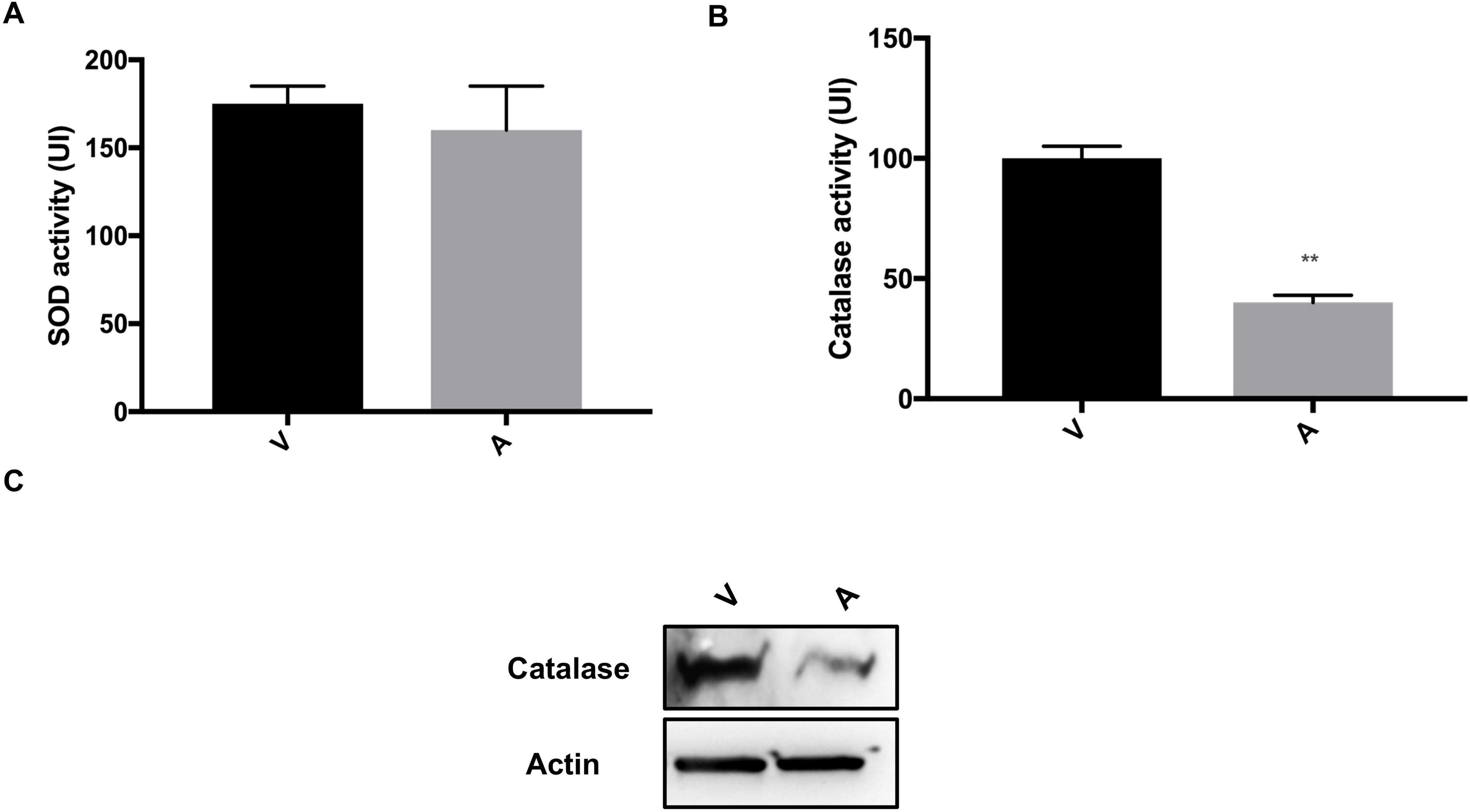
TGF-β2 activates *catalase* transcription in *Theileria*-transformed macrophages. **A.** The transcription of *catalase* and *GPx* is decreased in attenuated compared to virulent macrophages. Adding recombinant TGF-β2 restores transcription of *catalase* in attenuated macrophages to virulence levels. No effect was observed in the transcription level of *GPx*. **B.** Left panel, Catalase activity is down regulated in attenuated macrophages and rescued by exogenous TGF-β2 stimulation. **C.** Right panel, H_2_O_2_ levels are increased in attenuated compared to virulent macrophages and is dampened by addition of exogenous TGF-β2. All experiments were done independently (n = 3) and in triplicate. * p<0.05 compared to virulent macrophages; ** p<0.005 compared to virulent macrophages; *** p<0.001 compared to virulent macrophages; ## p<0.005 compared to attenuated macrophages and ### p<0.001 compared to attenuated macrophages.

### TGF-β2-signalling activates *catalase* transcription via CREB

CREB is a transcription factor that binds DNA on CRE sites (<u>c</u>AMP <u>r</u>esponse <u>e</u>lement) to regulate transcription of target genes, and in *Theileria*-transformed macrophages TGF-β2-signalling activates *CREB* transcription (8). Consistently, bioinformatic analyses detected CRE sites in the promoter region of the *catalase* gene (data not shown). Inhibition of CREB-mediated transcription with a specific CREB-CBP interaction inhibitor decreased *catalase* transcription and catalase activity to levels characteristic of attenuated macrophages (Figure 3A & B). Consequently, virulent macrophages produce H_2_O_2_ equivalent to attenuated levels and conversely, decreased H_2_O_2_ output occurred upon activation of CREB-mediated transcription following stimulation of attenuated macrophages with membrane-permeable db-cAMP (Fig. 3C). Importantly, pre-treatment of attenuated macrophages with the CREB inhibitor prevented the drop in H_2_O_2_ levels provoked by addition of db-cAMP (Fig. 3C). Thus, TGF-β2-signalling via CREB activates *catalase* transcription leading to increased catalase activity and reduced H_2_O_2_ output.

**Figure 3:**
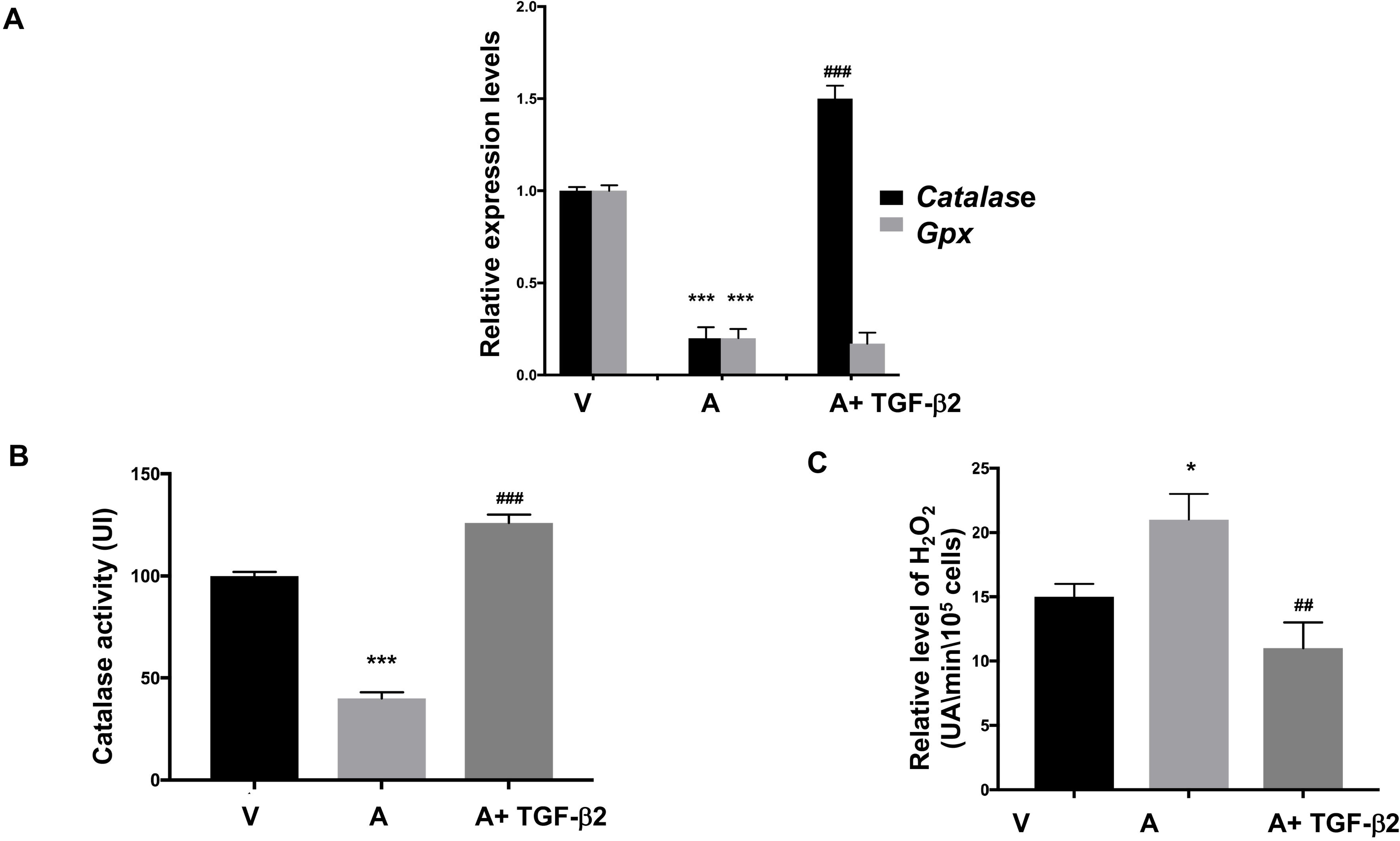
CREB drives *catalase* transcription. **A.** *Catalase* transcription is higher in virulent (V) than attenuated (A) macrophages. Catalase transcription is diminished to attenuated levels when CREB-mediated transcription in virulent macrophages is ablated with the specific CREB-CBP interaction inhibitor (25µM 1 h at 37°C). **B**. Catalase activity is higher in virulent (V) compared to attenuated (A) macrophages, and CREB-CBP-mediated inhibition of CREB in virulent macrophages reduces catalase activity to attenuated levels. **C.** H_2_O_2_ levels are higher in attenuated (A) compared to virulent (V) macrophages. CREB-CBP-mediated inhibition of CREB-driven transcription in virulent macrophages increases the level of H_2_O_2_, while adding db-cAMP to attenuated macrophages decreases H_2_O_2_ output. Treatment with CREBCBP interaction inhibitor abolishes *catalase* expression and ablates the increase in H_2_O_2_ induced by db-cAMP stimulation. All experiments were done independently (n = 3) and in triplicate. ** p<0.005 compared to virulent macrophages; *** p<0.001 compared to virulent macrophages and # p<0.05 compared to attenuated macrophages.

### Virulence and attenuation of *Theileria*-transformed macrophages depends on their redox balance

*T. annulata* transforms host macrophages into tumour-like cells that have heightened motility and invasiveness two traits that are typical of metastatic/disseminating cancer cells. Figure 4 shows that detoxifying H_2_O_2_ by adding catalase, or TGF-β2 to attenuated macrophages induces a regain in cell migration by *Theileria*-transformed macrophages. Boiling catalase to inactive the enzyme ablated its ability to reduce H_2_O_2_ levels that stimulate migration. By contrast, increasing H_2_O_2_ output by virulent macrophages via SB431542 blockade of TGF-R attenuated their migration (Fig. 4). Similarly, TGF-R blockade with SB431542 or inhibition of catalase activity with AminoTriazole (AT) reduced matrigel traversal of virulent macrophages to levels typical of attenuated macrophages (Fig. 4B). Therefore, modifying transformed macrophage redox balance via TGF-β2 stimulation, or manipulating catalase activity changes a virulence trait (heightened migration) of *Theileria*-transformed macrophages.

**Figure 4:**
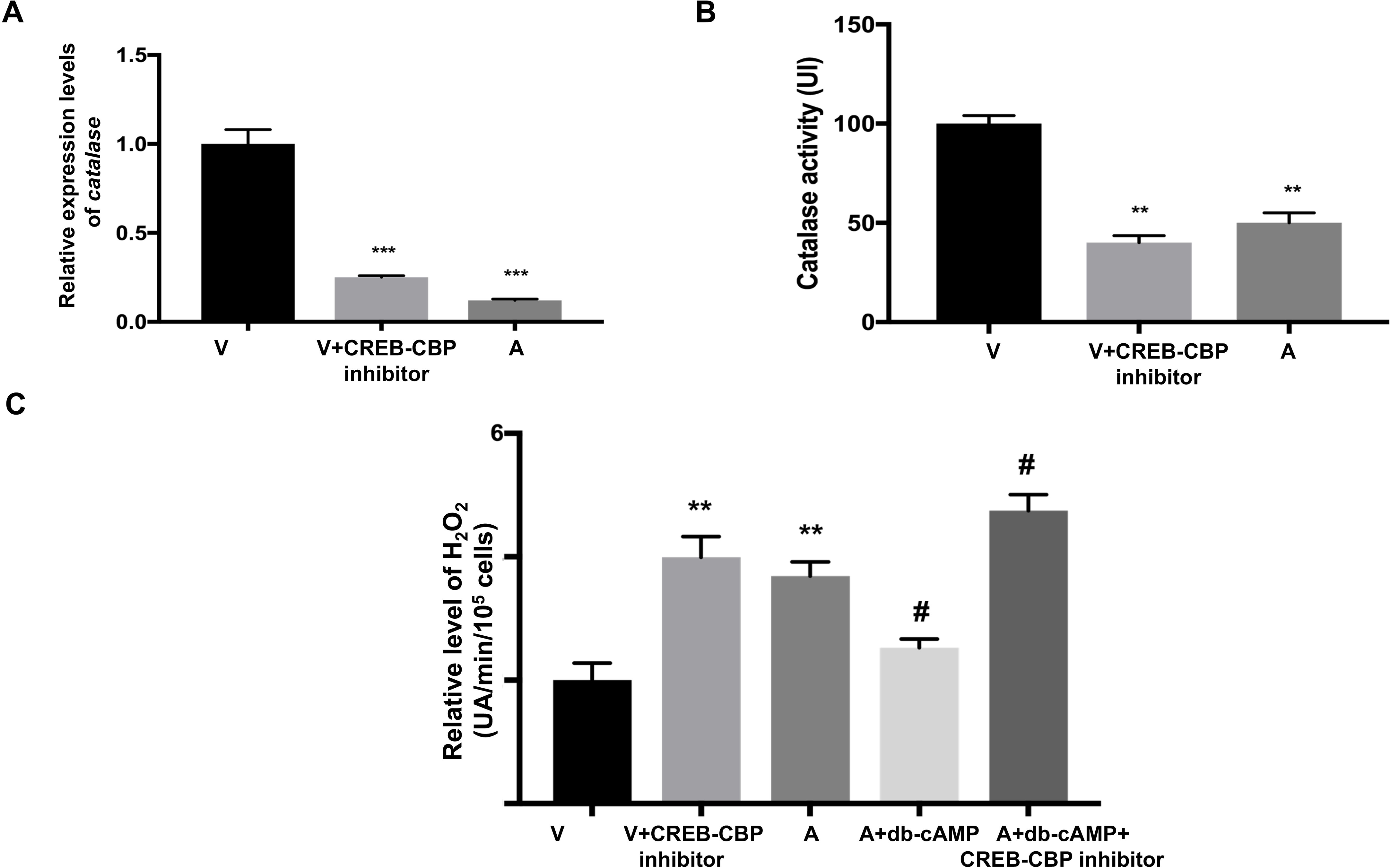
TGF-β levels regulate both oxidative stress and metastatic potential of *Theileria*-transformed macrophages. **A.** Cell migration index of virulent (V) macrophages is greater than that of attenuated (A) macrophages, but is reduced upon TGF-R blockade with SB431542. TGF-β2 stimulation of attenuated macrophages restores their cell migration index, as does addition of active catalase. Boiled catalase fails to restore attenuated macrophages migration index. **B.** Matrigel traversal is higher for virulent (V) than attenuated (A) macrophages. Blocking TGF-R-signalling with SB431542 diminishes traversal of V macrophages, as does AT-induced inhibition of catalase activity. All experiments were done independently (n = 3) and in triplicate. * p<0.05 compared to virulent macrophages; ** p<0.005 compared to virulent macrophages; *** p<0.001 compared to virulent macrophages; # p<0.05 compared to attenuated macrophages and # p<0.05 compared to attenuated macrophages.

### TGFβ-2 stimulation also increases catalase activity and metastatic potential of human lung and colon cancer cell lines

In order to extend our observations on *Theileria-*transformed macrophages to other cancer cell types, we treated HT-29 (human colorectal adenocarcinoma) and A549 (adenocarcinomic human alveolar basal epithelial) cell lines with TGF-R and/or CREB-CBP interaction inhibitors. Both HT-29 and A549 H_2_O_2_ levels increase following inhibitor treatment due to a corresponding drop in catalase activity (Figure 5 A& B), similar to *Theileria*-transformed macrophages (Figures 2, 3 & FigS1). Moreover, inhibition of TGF-R and/or CREB signalling decreases matrigel traversal of A549 cells (Fig. 5C). The ensembles lead to the conclusion that TGF-β2 regulation of catalase activity via CREB-mediated transcription and their impact on H_2_O_2_-type oxidative stress is common to different cancer cell types of human and bovine origin.

**Figure 5:**
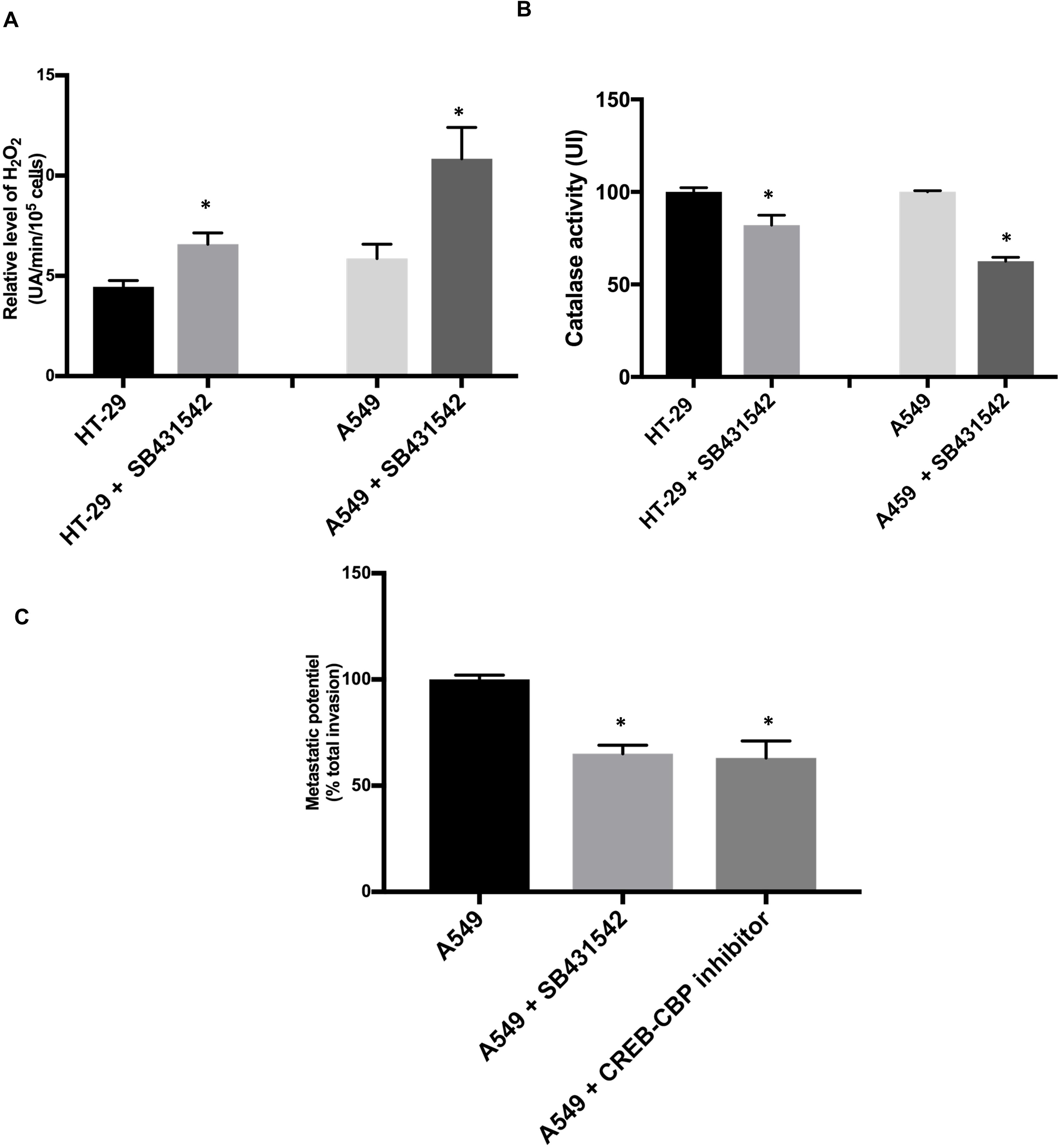
Observations on *Theileria*-transformed macrophages can be extended to human A549 and HT-29 cancer cell lines. **A.** H_2_O_2_ levels are lower in HT-29 and A549 compared to HT-29 and A549 treated with TGF-R inhibitor SB431542. **B**. Catalase activity is higher in HT-29 and A549 compared to HT-29 and A549 treated with SB431542. **C.** The metastatic potential as reflected by matrigel traversal of A549 decreased following CBP-induced inhibition of CREB and TGFR blockade by SB431542. * p<0.05 compared to HT-29 and HCT-116.

## Discussion

In this study, we have demonstrated that TGF-β2 induces CREB to drive *catalase* transcription, leading to more catalase enzyme and hence activity to detoxify H_2_O_2_. *Theileria*-transformed macrophages with attenuated dissemination potential are characterized by decreased TGF-β2 production, and consequently, reduced catalase activity and increased H_2_O_2_ output. Stimulating attenuated macrophages with exogenous TGF-β2 increased catalase activity and decreased H_2_O_2_ output leading to a regain in their capacity to migrate and traverse matrigel. In contrast, blockade of either TGF-R or CREB binding to CBP decreased catalase activity and increased H_2_O_2_ output led to a reduced migratory and matrigel traversal capacity of virulent macrophages. These observations are not restricted to *Theileria*-transformed bovine macrophages, but were shared by human A549 and HT-29 cancer cell lines, where inhibition of TGF-β2-signalling and/or CREB-mediated transcription again decreased catalase activity, increased the H_2_O_2_ output and reduced their capacity to traverse matrigel. Thus, our demonstration that catalase activity and hence H_2_O_2_ levels are regulated by TGF-β2-signaling can be generalized to different types of tumours.

Our study is consistent with a pro-metastatic role for TGF-β2, since adding back recombinant TGF-β2 to attenuated macrophages resulted in a regain in their migratory and dissemination potentials. Clearly, one way that TGF-β2 promotes virulent dissemination is by inducing CREB transactivation to activate catalase and detoxify excess H_2_O_2_. Tumours produce large amounts of ROS that cause damage to DNA, proteins and lipids and we propose that infected macrophages attenuated for dissemination have countered the tumorigenic effect of *Theileria* by producing higher levels of H_2_O_2_. In virulent macrophages with high TGF-β2 levels CREB induction activates catalase that dampens H_2_O_2_ output and similarly TGF-β2-mediated changes H_2_O_2_ output and catalase activity impact on the metastatic potential of human A549 and HT-29 cancer cell lines. This argues that many tumours of different origins exploit TGF-β2-driven *catalase* expression to control their H_2_O_2_ redox balance.

## MATERIALS AND METHODS

### Cell culture

virulent Ode macrophage line (12) corresponds to passage 62 and attenuated macrophages correspond to passage 309. All macrophages were maintained in RPMI medium supplemented with 10% fetal bovine serum (FBS), 2 mM L-glutamine, 100U penicillin, 0.1 mg/ml streptomycin, and HEPES. The cell lines S2, S3 and H7–H8 have been described previously (13). Cells were maintained in the above culture medium with 50 µM of β-mercaptoethanol. The human colon adenocarcinoma cell line HT-29 was maintained in McCoy’s medium supplemented with 10% FBS. A549 human lung adenocarcinoma cells (ATCC, CCL-185) were cultured in DMEM/RPMI (1:1) with 10% fetal bovine serum. All the cells were incubated at 37°C with 5% CO_2_.

### Total RNA extraction and reverse transcription

Total RNA of *Theileria*-infected macrophages was isolated using the RNeasy mini kit (Qiagen) according to the manufacturer’s instructions. The quality and quantity of RNA was measured by Nanodrop spectrophotometer. For reverse transcription, 1µg isolated RNA was diluted in water to a final volume of 12µL, warmed at 65°C for 10 min, then incubated on ice for 2 min. Afterwards, 8µl of reaction solution (0.5µL random hexamer, 4µL 5x RT buffer, 1.5µL 10mM dNTP, 1µL 200U/µLRT-MMLV (Promega) and 1µL 40U/µLRNase inhibitor (Promega) was added to get a final reaction volume of 20µL and incubated at 37°C for 2 h. The resultant cDNA was stored at −20°C.

### Quantitative polymerase chain reaction (qPCR)

mRNA expression levels were estimated by qPCR on Light Cycler 480 (Roche) using SYBR Green detection (Thermo). *GAPDH* was used as internal control to normalize for mRNA levels. The detection of a single product was verified by dissociation curve analysis and relative quantities of mRNA calculated using the method described (14).

### Pharmaceutical inhibition and activation

TGF-β signalling was inhibited using the TGF-Receptor I/ALK5 inhibitor SB431542, 10µM (Sigma #S4317) and activated by adding 5ng/ml of recombinant bovine TGF-β2 (NIBSC, Potters Bar. UK). Cells were treated for 24 h at 37°C. Catalase activity was ablated with a selective inhibitor Aminotriazole (AT) (Sigma, A8056), and restored by adding bovine catalase (Sigma, C4963-2MG). Cells were treated overnight at a concentration of 1200µM with AT and 80U/ml of bovine catalase. For inhibiting CREB transcription, a cell-permeable naphthamide compound that effectively blocks the interaction between the KIX domain of CBP and the KID domain of CREB was used (Calbiochem, CAS 92-78-4) at a concentration of 25µM for 1 h at 37°C with 5% CO_2_.

### Western Blotting

Cells were harvested and extracted by lysis buffer (20mM Hepes, Nonidet P40 (NP40) 1%, 0.1% SDS, 150mM NaCl, 2mM EDTA, phosphatase inhibitor cocktail tablet (PhosSTOP, Roche) and protease inhibitor cocktail tablet (Complete mini EDTA free, Roche). Protein concentration was determined by the Bradford protein assay. Cell lysates were subjected to Western blot analysis using conventional SDS/PAGE and protein transfer to nitrocellulose filters (Protran, Whatman). The membrane was blocked by 5% non-fat milk-TBST (for anti-catalase), or 3% non-fat milk-PBST (for anti-actin antibody) for 90 min at room temperature (RT).

Antibodies used in immunoblotting were as follows: rabbit polyclonal antibody anti-catalase (Cell Signaling) and goat polyclonal antibody anti-actin (I-19, Santa Cruz Biotechnology). Membranes were incubated with peroxidase-conjugated secondary antibody (rabbit anti-IgG and goat anti-IgG (Santa Cruz biotechnology). After washing, proteins were visualized with ECL western blotting detection reagents (Thermo Scientific) on X-ray films. The level of β-actin was used as a loading control throughout.

### Dynamic monitoring of cell migration with the xCELLigence system

Cell migration assay was assessed using the xCELLigence system (Roche). Medium was added to the bottom chamber of the CIM-Plate 16. The CIM-Plate 16 was assembled by placing the top chamber coated with Matrigel (BD) onto the bottom chamber and snapping the two together. Serum-free medium was placed in the top chamber to hydrate and the membrane was pre-incubated for 1 hour in the CO2 incubator at 37°C. Once the CIM-Plate 16 has equilibrated, it is placed in the RTCA DP station and the background cell-index values are measured. The CIM-Plate 16 is then removed from the RTCA DP station and the cells passaged 24h in serum free medium were added to the top chamber. The CIM-Plate 16 is placed in the RTCA DP station and migration is monitored for several hours.

### Matrigel chambers assay

The invasive capacity of transformed cells was assessed *in vitro* using Matrigel migration chambers, as described (15). Culture coat 96-well medium BME cell invasion assay was obtained from Culturex instructions (3482-096-K). After 24 h of incubation at 37°C, each well of the top chamber was washed once in buffer. The top chamber was placed back on the receiver plate. 100µL of cell dissociation solution/Calcein AM were added to the bottom chamber of each well, incubated at 37°C for 1 h to fluorescently label cells and dissociate them from the membrane before reading at 485 nm excitation, 520 nm emission using the same parameters as the standard curve.

### Intracellular levels of hydrogen peroxide (H_2_O_2_)

1×10^5^ cells were seeded in 96 well plates and incubated 18h in complete medium. Cells were washed in PBS and incubated with 100μL per well of 5μM H2-DCFDA diluted in PBS (Molecular Probes). H_2_O_2_ levels were assayed by spectrofluorimetry on a fusion spectrofluorimeter (PackardBell). Fluorescence intensity was recorded every hour over a period of 5 h. Excitation and emission wavelengths used for H_2_O_2_ were 485 and 530nm. The number of cells was evaluated by the crystal violet assay. Cells were stained in 0.05% crystal violet and 2% ethanol in PBS for 30 min at room temperature. After four washes in PBS, the stain was dissolved in methanol and measured at 550nm on Fusion. The level of H_2_O_2_ was calculated in each sample as follows: reactive oxygen species rate (arbitrary unitsmin^−1^10^5^ cells^−1^) = [fluorescence intensity (arbitrary units) at T300 minutes − fluorescence intensity (arbitrary units) at T0] per 60min per number of cells as measured by the crystal violet assay.

### Catalase activity assay

A dry pellet of 1×10^5^ cells was lysed in 50μL PBS, 1% NP40. 50μL of lysate, 50μL of anti-peroxydase antibody (1/2000, Sigma) and 50μL H_2_O_2_ (1/4000, Sigma) were added to a 96-well plate and incubated for 10 min at 37°C in 5% CO_2_. 50μL of OPD (SIGMAFAST^TM^, #P9187) was then added and the absorbance immediately read at 405nm. Catalase activity assay was assayed on Fusion. Catalase measurement was reported to the amount of protein in each sample (bovine serum albumin microbiuret assay, Pierce, Bezons, France).

### SOD activity

Superoxide dismutase (SOD) activities of cells were evaluated by the nitroblue tetrazolium reduction technique according to Beauchamp and Fridovich (16). SOD measurements were reported to the amount of protein in each sample (Bradford method was used).

### Statistical Analysis

Data were analyzed with the Student’s t-test. All values are expressed as mean+/-SEM. Values were considered to be significantly different when p values were < 0.05.

## Acknowledgements

We thank Arnab Pain for fruitful discussion and input when writing the manuscript. MH was supported by a PhD CNR fellowship from the Lebanese government. GL acknowledges support from ANR grant (11 BSV3 01602), Labex ParaFrap (ANR-11-LABX-0024) and core support from INSERM and the CNRS.

---

**Figure S1:**
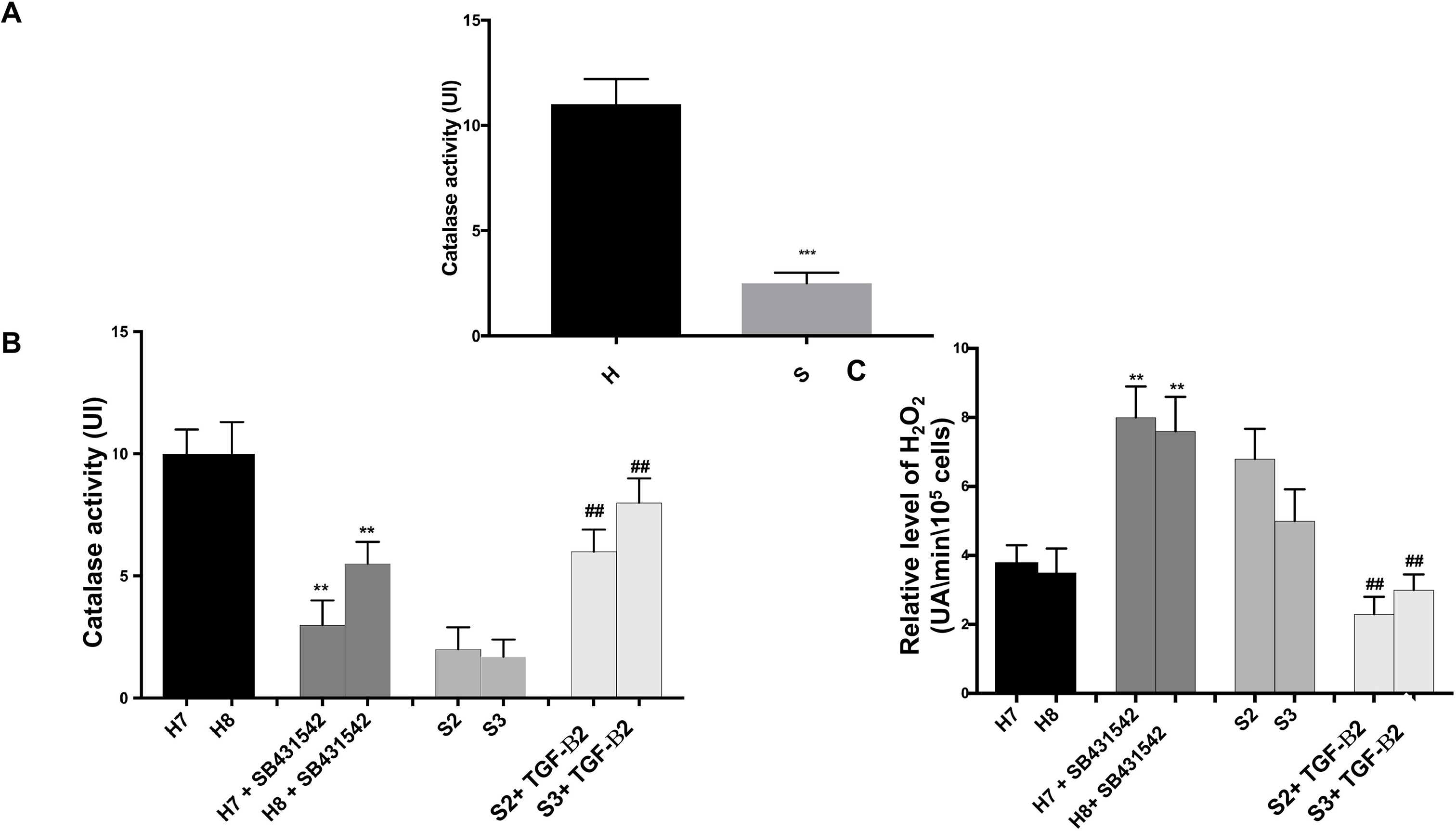
TGF-β2 regulates oxidative stress in *Theileria*-transformed macrophages. **A and B.** Catalase activity is augmented in independent clones (H7 & H8) of macrophages isolated from disease-susceptible Holstein-Friesian (H) animals compared to independent clones (S2 &S3) of macrophages isolated from disease-resistant Sahiwal (S) animals. SB431542 blockade of TGF-R abolished heightened catalase expression by independent clonal lines of H macrophages, whereas adding TGF-β2 to independent clonal lines of S macrophages increased catalase activity. **C.** H_2_O_2_ output is higher in disease-resistant S macrophages compared to disease-susceptible H macrophages. Blockade of TGF-R-signalling in H macrophages increased levels H_2_O_2_, while stimulating independent clonal lines of S macrophages with TGF-β2 decreased H_2_O_2_ levels. ** p<0.005 compared to Holstein macrophages and ## p<0.05 compared to Sahiwal macrophages.

